# RSPO2-based peptibodies conjugated with pyrrolobenzodiazepine dimer or camptothecin analogs demonstrate potent anti-tumor activity by targeting the three receptors LGR4/5/6 in colorectal cancer and neuroblastoma

**DOI:** 10.1101/2025.04.25.650662

**Authors:** Yukimatsu Toh, Jianghua Tu, Ling Wu, Adela M. Aldana, Jake J. Wen, Lynn H. Su, Bin Yang, Xiaowen Liang, Li Li, Sheng Pan, Jin Wang, Jie Cui, Qingyun J. Liu

## Abstract

Leucine-rich repeat containing, G protein-coupled receptor 4, 5, and 6 (LGR4/5/6) are three homologous receptors that are co-expressed or alternately expressed at high levels in tumor cells of colorectal cancer (CRC) and high-risk neuroblastoma (NB). Simultaneous targeting of all three receptors may provide increased efficacy or overcome drug resistance due to tumor heterogeneity and cancer cell plasticity. LGR4/5/6 all bind to R-spondins (RSPOs) with high affinity and potentiate Wnt/β-catenin signaling in response. Previously, we showed that a peptibody based on a mutant RSPO4 furin domain that bound to LGR4/5/6 without potentiating Wnt/β-catenin signaling was able to deliver cytotoxins into cancer cells that express any of the three receptors. We have now generated a mutant RSPO2 furin domain that retains high affinity binding to LGR4/5/6 without signaling activity. Peptibodies based on this RSPO2 furin mutant were conjugated with either pyrrolobenzodiazepine dimer (PBD) or camptothecin derivative (CPT2), and the resulting peptibody-drug conjugates (PDCs) showed potent and specific cytotoxic activity in NB and CRC cell lines expressing any of LGR4/5/6 *in vitro* and robust anti-tumor activity *in vivo*. The results support the potential of RSPO2-based PDCs for the treatment of CRC, high-risk NB, and other cancers that express any of LGR4/5/6.

## INTRODUCTION

Colorectal cancer (CRC) and high-risk neuroblastoma (NB) remain major clinical challenges with limited durable responses under current standard-of-care therapies. In metastatic CRC, frontline treatment typically consists fluoropyrimidine-ased chemotherapy combined with oxaliplatin or irinotecan, often paired with targeted agents such as anti-VEGF or anti-EGFR antibodies depending on RAS mutation status^1^. More recently, immune checkpoint inhibitors have provided substantial benefit in microsatellite-instable/mismatch repair deficient (MSI-H/dMMR) CRC, yet most microsatellite-stable tumors remain refractory to immunotherapy^1^. High-risk NB is treated with multi-agent chemotherapy, anti-GD2 immunotherapy, radiotherapy, and autologous stem-cell rescue, yet relapse rates remain high and effective salvage options are limited^2, 3^. These clinical gaps underscore the need for new targeted modalities, including antibody–drug conjugates (ADCs), peptide- or ligand-directed targeting therapeutics, and engineered biologics designed to exploit tumor-specific receptor expression^4^.

Leucine-rich repeat containing, G protein-coupled receptor 4, 5, and 6 (LGR4/5/6) are three related membrane receptors with a large extracellular domain (ECD) and a seven transmembrane (7TM) domain typical of the rhodopsin family of G protein-coupled receptors ^5-7^. LGR4 is broadly expressed at low levels in epithelial tissues with critical roles in cell proliferation and migration during organ development whereas LGR5 is expressed mostly in adult stem cells in the gastrointestinal (GI) tract^6, 8, 9^. R-spondins are a group of four related secreted proteins (RSPO1/2/3/4) that play essential roles in normal development and survival of adult stem cells^10^. RSPOs bind to LGR4/5/6 with high affinity and potentiate canonical Wnt/β-catenin signaling^11-14^. Mechanistically, RSPO and LGR4 form a complex to inhibit the function of RNF43 and ZNRF3, two E3 ligases that ubiquitinate Wnt receptors for degradation following Wnt-ligand induced receptor activation, leading to sustained and stronger signaling^15-18^. In contrast, LGR5 does not potentiate Wnt signaling through sequestering of the E3 ligases^17^.

LGR4/5/6 are often co-expressed or alternately expressed in various cancer types, particularly in cancers of the gastrointestinal system, with LGR4 co-expressed with LGR5 or LGR6^19-24^. Furthermore, LGR5 has been shown to be enriched in cancer stem cells of colorectal cancer (CRC)^25-28^ and LGR5-positive cancer cells were shown to fuel the growth of primary tumors and metastasis^29, 30^. The Cancer Genome Atlas’s (TCGA’s) RNA-Seq data of colorectum, liver, and stomach cancers confirmed that LGR4 expression at high levels in nearly all cases while LGR5 and LGR6 were co-expressed with LGR4 in the majority of CRC and substantial fractions of liver and stomach cancer. LGR4/5/6 were also found to be highly expressed in liver metastasis of CRC^31, 32^. On the other hand, LGR5 was found to be one of the genes that were most enriched in NB cells selected for sphere-forming and metastasis capability^33^. Subsequent studies reported that LGR5 expression is associated with aggressive diseases in NBs with or without *MYCN* amplification^34-37^. LGR5 expression was also found to be highly enriched in end-stage tumors in the mouse model of NB driven by *MYCN* overexpression^38^. Knockdown of LGR5 in LGR5-high NB cell lines led to reduced MAPK signaling and decreased cell growth^35^. Recently, LGR5 expression was found to be increased in drug-resistant NB cell lines^39^. Overall, these expression data support that simultaneous targeting of LGR4/5/6 may provide an effective approach for the treatment of metastatic CRC and high-risk NB.

Antibody-drug conjugates (ADCs) have become a major modality of cancer therapy^4, 40^. ADCs consist of a monoclonal antibody linked to a cytotoxic payload via a chemical linker^4, 40^. The antibody component specifically targets antigens overexpressed on cancer cells, delivering potent cytotoxic agent directly to the tumor, thereby reducing systemic toxicity. Recent progress in linker chemistry have further improved efficacy with reduced toxicity^41, 42^. Previously, we reported that anti-LGR5 ADC conjugated with pyrrolobenzodiazepine dimer (PBD) had robust anti-tumor activity in NB tumor models^37^. We also reported the construction of an RSPO4 peptibody, which consists of a human IgG1-Fc conjugated with the cytotoxin monomethyl auristatin E (MMAE) or a camptothecin analog (CPT2) while fused to a modified RSPO4 furin domain. This peptibody can then deliver the drugs into any LGR4/5/6-expressing cancer cells^43, 44^. This delivery is similar to the ADC approach except the ligand-drug conjugate enables the targeting of LGR4/5/6 simultaneously. Here we report the generation and characterization of RSPO2-based peptibody-drug conjugate (PDC) using PBD and CPT2 in preclinical models of CRC and NB. The PBD- and CPT2-based RSPO2 PDCs showed highly potent cytotoxic activity across LGR4/5/6-expressing CRC and NB cell lines *in vitro* and robust anti-tumor effect in xenograft models *in vivo*.

## RESULTS AND DISCUSSION

### An RSPO2 furin domain mutant that retains high affinity binding to LGR4/5 without potentiating Wnt/β-catenin signaling was identified

Of the four RSPOs, RSPO4 furin domain has the highest affinity to LGR4/5/6 with RSPO2 being a close second^45, 46^. RSPO2 is also unique for its high affinity binding to RNF43/ZNRF3 in the absence of LGR4/5/6 ^46, 47^. We then evaluated the potential use of RSPO2 furin domain peptibody for drug conjugation. X-ray crystal structure data showed that M68 and Q70 of RSPO2 furin-1 domain are critical to binding to RNF43/ZNRF3 and thus essential for potentiating of Wnt/β-catenin signaling ^47, 48^ (Fig. 1A). Previously, we fused wild-type RSPO2 furin domain to human IgG1-Fc domain to create a peptibody designated R203 (Fig. 1B). We first generated a Q70R mutant called R206, similar to that of RSPO4. Unlike RSPO4, mutation of Q70R alone was only able to partially inactivate W/β-catenin signaling^47^ (Fig. 1D). We then evaluated various M68 mutations in combination with Q70R and found that the M68T/Q70R double mutation (designated R232, Fig. 1B) resulted in nearly complete loss of activity in potentiating Wnt/β-catenin signaling (Fig. 1C-D). In binding analysis, R232 showed high affinity binding to cells expressing either LGR4 (Kd = 0.2 nM) or LGR5 (Kd = 0.3 nM) (Fig. 1E), similar to that of wild-type RSPO2 peptibody R203^46^, with little binding to vector control cells (Fig. 1E).

**Figure 1.**
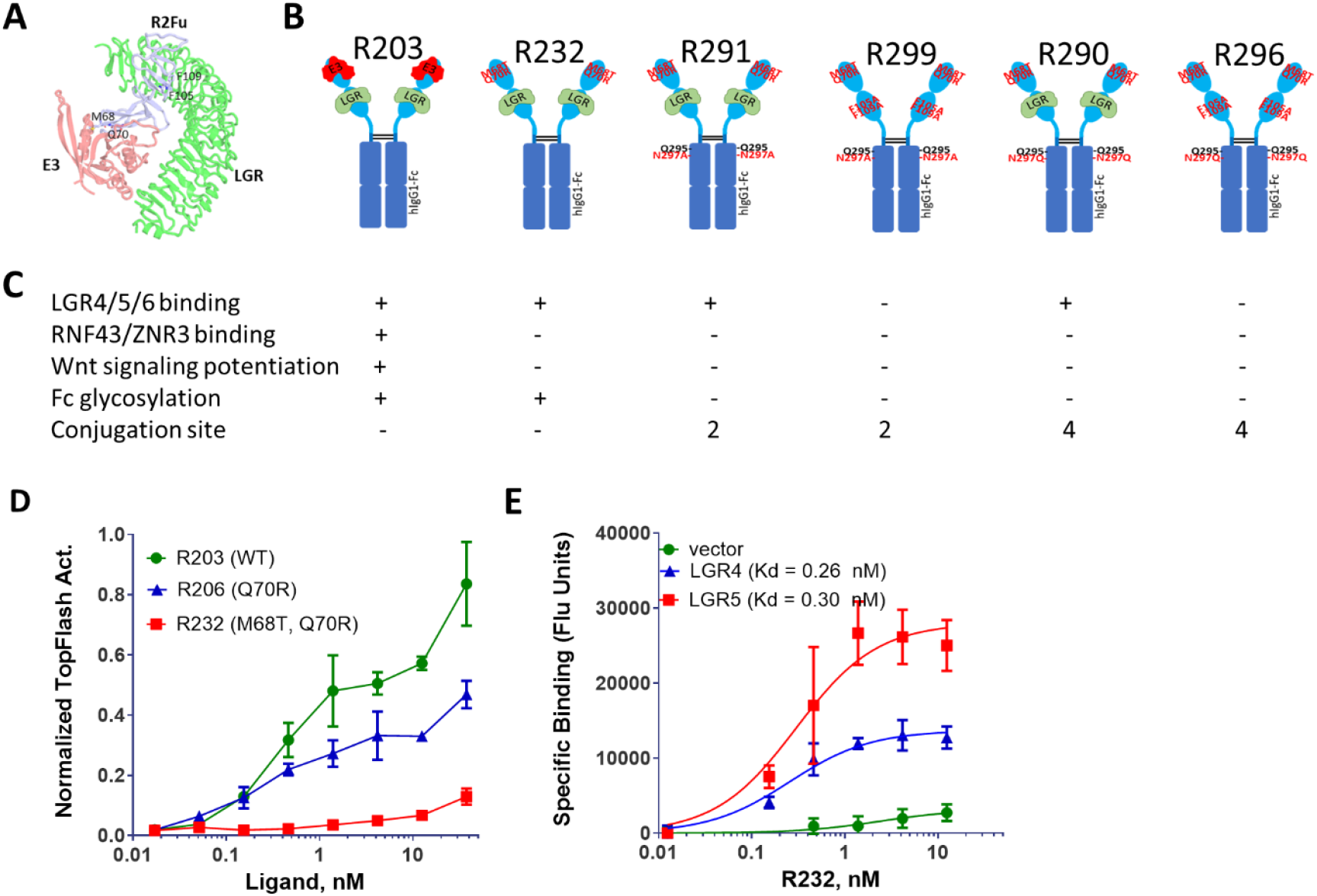
Generation of RSPO2-furin Fc fusion proteins and characterization of their binding affinities. A, Structure model of RSPO2 furin domain bound to E3 ligase and LGR4ECD. B, Schematic diagram of RSPO2 furin domainbased peptibodies with various mutations. C, Summary of binding and signaling activity of the various RSPO2 furin peptibodies. D, Potentiation of Wnt/p-catenin signaling by R203, R206, and R232 in the TopFlash assay. E, Whole-cell binding of R232 to HEK293T cells stably expressing LGR4, LGR5 or vector control. Error bars are SEM (N = 2-3).

For site-specific conjugation of linker-payload, we introduced an N297A change into the glycosylation site of the Fc domain to create the peptibody designated R291, which has a glutamine acceptor site (Q295) for the microbial transglutaminase^49^ (Fig. 1B). For a control peptibody, we generated R299 which has two additional mutations in RSPO2 furin-2 domain (F105A/F109A) that abolishes binding to LGR4/5/6 as predicted by crystal structure data of RSPO2 binding to LGR5^48^ (Fig. 1B-C, Fig. 2A). Binding analysis confirmed that R291 had high affinity binding to LGR4/5/6 whereas R299 had much reduced binding (Fig. 2D-F). To evaluate the potential of RSPO2 furin domain peptibodies for drug delivery, we conjugated the potent, PBD-based cytotoxin SG3199 to R291 and R299 using a chemoenzymatic method^37, 49, 50^ (Fig. 2A). An amino-PEG4-azide was first attached to R291/R299 with microbial transglutaminase and complete conjugation was confirmed by mass spectrometry. DBCO-PEG8-VA-PAB-SG3199 was then linked to the azide through click chemistry and the resulting PDCs (R291-SG3199 and R299-SG3199) showed complete conjugation, which were confirmed to have two payloads per peptibody using mass spectrometry (Fig. 2B-C). Binding analysis showed that R291-SG3199, similar to unconjugated R291, retained high affinity binding to cells expressing LGR4, LGR5, or LGR6 whereas R299-SG3199 had much lower affinity binding (Fig. 2G-I).

**Figure 2.**
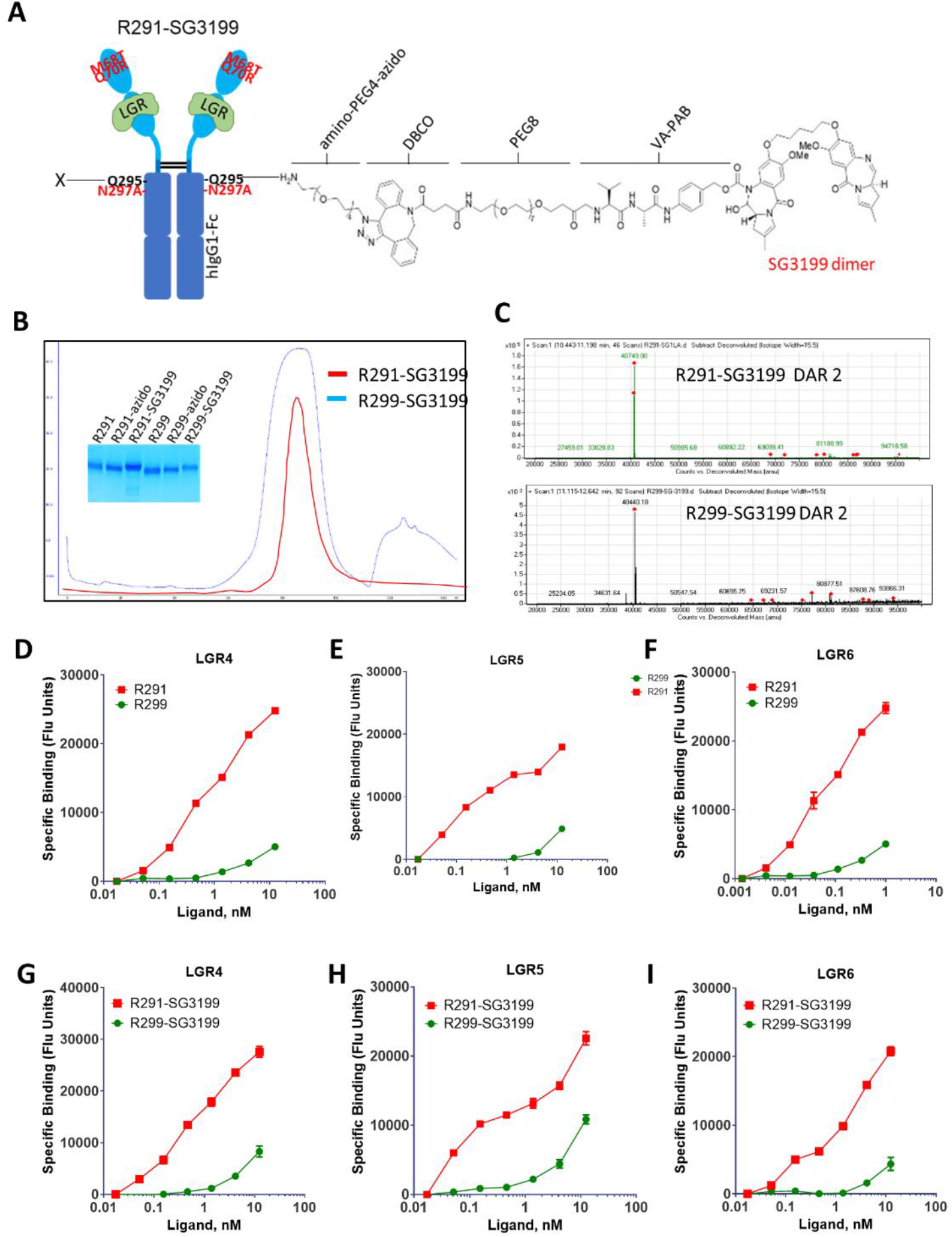
R291 and R299 PDC synthesis and conjugation A, Schematic diagram of R291/R299 conjugated with SG3199. B. Chromatogram of R291- and R299-SG3199 in gel filtration and Coomassie staining of SDS-PAGE of R291/R299 before and after conjugation. C, Mass spectrum of R291-SG3199 (top) and R299-SG3199 (bottom). D-F, Whole-cell binding of unconjugated R291 and R299 to HEK293T cells expressing LGR4 (D). LGR5 (E). and LGR6 (F). O-l, Whole-cell binding of R291-SG3199 and R299-SG3199 to HEK293T cells expressing LGR4 (G), LGR5 (H), and LGR6 (I). Error bars are SEM (N = 2-3).

### Cytotoxicity of Free Payloads in Neuroblastoma Cell Lines

To establish the baseline cytotoxicity of the individual payloads, we evaluated the cytotoxicity of the corresponding free drugs—MMAE, the PBD dimer SG3199, the topoisomerase-I inhibitor deruxtecan (DXd), and a topoisomerase I inhibitor camptothecin analog—SN38 in a panel of neuroblastoma cell lines (SK-N-AS, SK-N-BE2, CHP212, and CHP212-LGR5KO). As shown in Supplementary Figure S1, each payload exhibited a dose–dependent cytotoxicity. Free PBD displayed the highest potency across all NB cell lines, with sub-nanomolar IC_50_ values, whereas SN38 showed only modest activity, yielding IC_50_ values in the low-to mid-nanomolar range. DXd and MMAE demonstrated intermediate potencies, with IC_50_ values spanning approximately 0.1–1 nM depending on the cell line. Importantly, CHP212-LGR5KO cells exhibited similar sensitivity to the free payloads as the parental CHP212 line, confirming that the intrinsic cytotoxicity of each small-molecule drug is independent of LGR expression. These results establish a baseline potency ranking and provide a benchmark for interpreting the enhanced selectivity and therapeutic index achieved by the RSPO2-based PDCs relative to the unconjugated drugs.

### RSPO2 furin mutant peptibody conjugated with PBD had potent and specific cytotoxic activity against cancer cells expressing LGR4/5/6 *in vitro*

R291-SG3199 and R299-SG3199 were tested side-by-side in a panel of NB cell lines that express LGR5 at high levels as we showed previously^17, 37^. In CHP212 cells, R291-SG3199 showed an IC50 of 0.09 nM whereas the control PDC R299-SG3199 showed an IC50 of 4.9 nM (Fig. 3A). In CHP212-LGR5KO cells,in which LGR5 was knocked out by CRISPR-Cas9 as we described before^37^, both R291-SG3199 and R299-SG3199 displayed similar potency with an IC50 of approximately 10 nM (Fig. 3A). In two other NB cells expressing LGR5 (SKNBE2 and SKNAS)^35, 37^, R291-SG3199 also showed potent cytotoxic activity (IC50 = 0.01 nM) whereas R299-SG3199 was much less potent (Fig. 3B). These results indicate that R291-SG3199 has a potent and specific cytotoxic activity in NB cells expressing LGR5.

**Figure 3.**
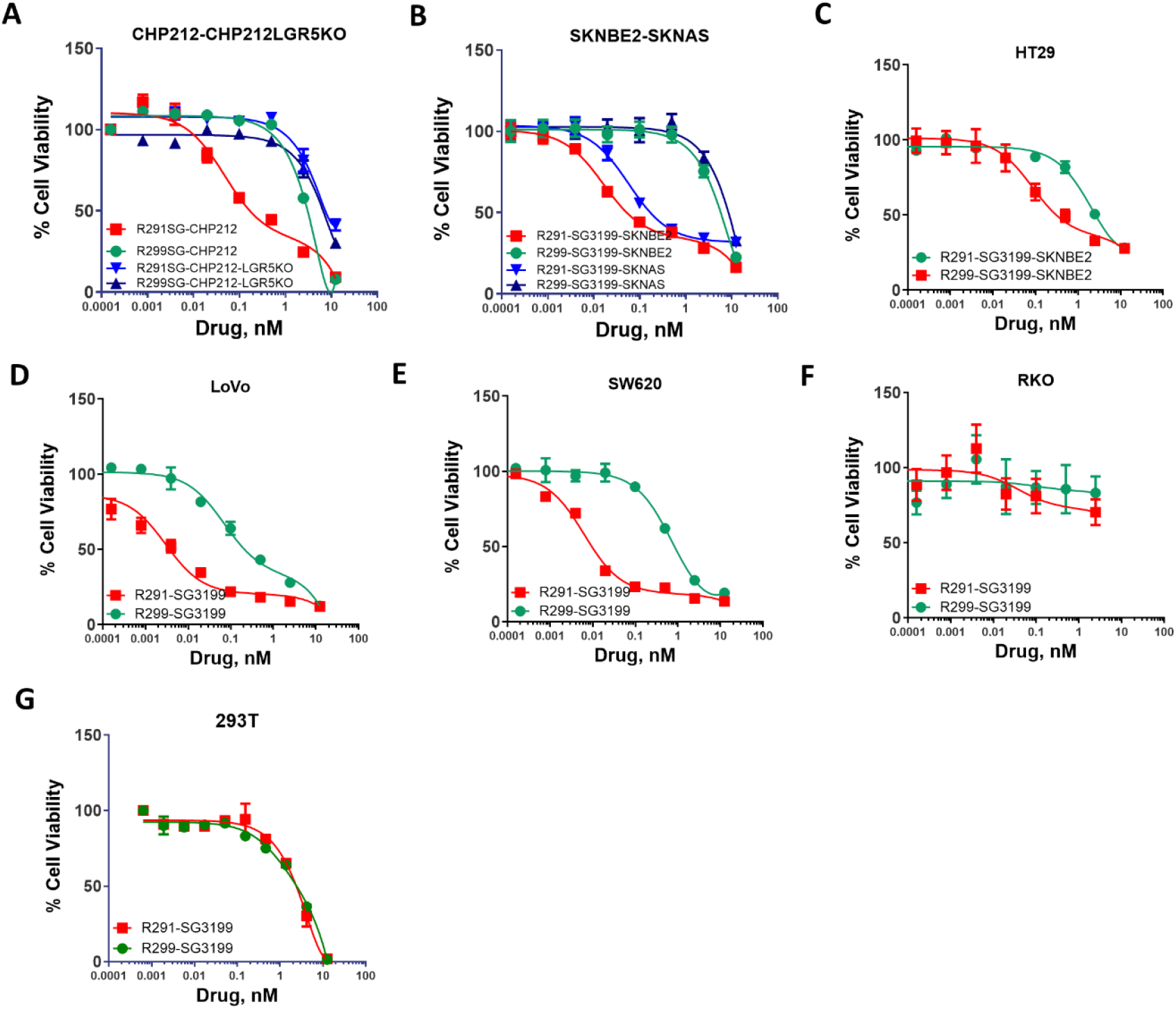
Cytotoxic activity of R291-SG3199 and R299-SG3199 in NB and CRC cell lines.. A-B, NB cell limes CHP212 and CHP212-LGR5KO (A) and SKNBE2 and SKNAS (B). C-F, CRC cell lines HT29 (C). LOVO (D). SW620 (E). and RKO (F). G, HEK-293T cell line. Error bars are SEM (N = 2-3).

We then tested the two PDCs in a panel of colorectal cancer cells expressing LGR4, LGR5 and/ or LGR6 both as we had described^43^. In HT29 cells, which express only LGR4, R291-SG3199 has an IC50 of 0.1 nM while R299-SG3199 has an IC50 of 5 nM (Fig. 3C). In LoVo and SW620 cells, which express all three receptors at various levels^43^, R291-SG3199 had an IC50 of ∼0.01 nM whereas R299-SG3199 displayed an IC50 of ∼1 nM (Fig. 3D-E). In RKO cells, which express none of the three LGRs according to CCLE’s RNA-seq data, R291-SG3199 and R299-SG3199 showed no cytotoxic activity at up to 3 nM (Fig. 3F). In HEK293T cells which are normal kidney epithelial cells, R291-SG3199 and R299-SG3199 also had similar potency with IC50 of 3.0 nM. These results showed that R291-SG3199 is ∼10-100x more potent in cancer cell lines expressing any of LGR4/5/6 whereas the two PDCs had similar potency in cells lacking expression of any of the three receptors, indicating that the cytotoxic activity of R291-SG3199 was mostly specifically mediated by LGR4/5/6. The cytotoxic activity of the control PDC R299-SG3199, though being much less potent than R291-SG3199, is most likely a result of non-specific uptake by the cancer cells such as pinocytosis or residual binding activity to LGR4/5/6.

### R291-SG3199 had robust anti-tumor activity in NB xenograft models *in vivo*

To evaluate the activity of RSPO2-based PDC *in vivo*, we tested R291-SG3199 in xenograft models of NB cell lines, SKNAS and SKNBE2. First, we tested the stability of R291 and R291-SG3199 in mice by determining its concentration in blood following dosing at 1 mg/kg by intraperitoneal injection. Both conjugated and unconjugated R291 only had a half-life of ∼6 hours, indicating rapid clearance (Supplementary Fig. S2). For the SKNAS model, R291-SG3199 was given at 0.15 mg/kg and 0.3 mg/kg, twice per week for 4 weeks, along with the vehicle control. As shown in Figure. 4A, R291-SG3199-treated mice had reduction in tumor growth, with the low dose (0.15 mg/kg) reducing tumor growth by 45% (p = 0.12 vs vehicle, One-Way ANOVA) and the high dose (0.3 mg/kg) reducing tumor growth by 87% (p = 0.002 vs vehicle). No major adverse effect and no significant difference in body weights among the groups were observed, suggesting that the PDC was tolerated at these dose levels (Fig. 4B). Importantly, both R291-SG3199 groups extended survival significantly, with median survival of 39 days for the low dose group vs. 25 days for the vehicle group (p = 0.001, log-rank test), and 46 days for the high dose group (p < 0.0001 vs. vehicle, log-rank test) (Fig. 4C). We then evaluated R291-SG3199 in xenograft models of the NB cell line SKNBE2. The PDC was given at 0.3 mg/kg, twice per week for a total of eight doses. As shown in Figure. 4D, R291-SG3199 showed significant inhibition of tumor growth, with a 73% reduction (p = 0.03 vs vehicle control, T-test) in tumor growth as measured on Day 25 (the last day before the vehicle group had animals sacrificed). PDC was tolerated at these dose levels as no major adverse effect was observed and no significant difference found in body weights among the groups was noted (Fig. 4E). Similarly, the PDC also extended survival significantly, with median survival of 61 days vs 25 days for the vehicle control group (p < 0.0001, log rank-test) (Fig. 4F). These *in vivo* results demonstrate that R291-SG3199 was able to significantly inhibit tumor growth at 0.3 mg/kg, which is approximately twice the human dose level of the SG3199-based ADC loncastuximab tesirine^50^.

**Figure 4.**
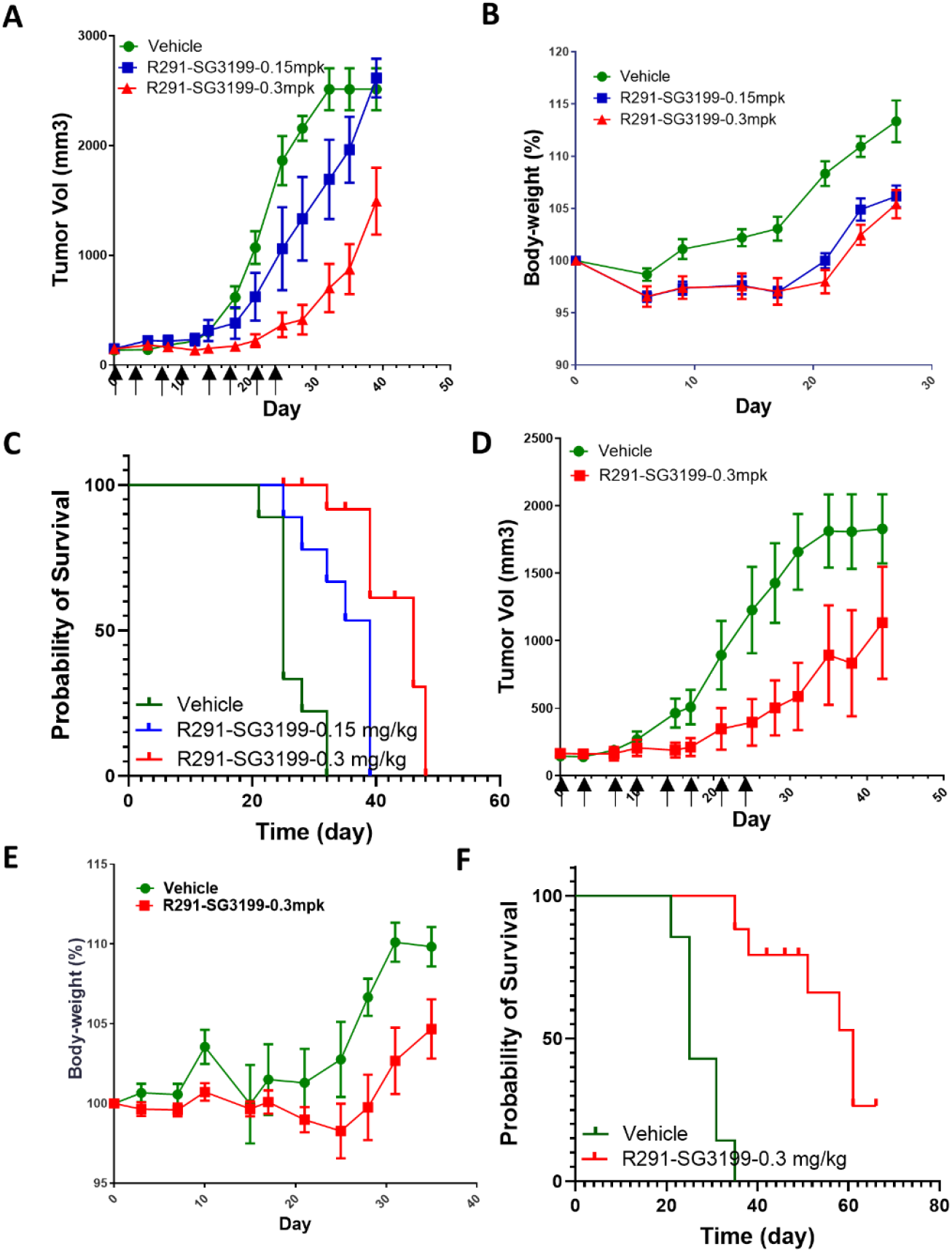
Anti-tumor activity of R291-SG3199in xenograft models of NB cell lines. A, Tumor growth curves of SKNAS xenografts treated with vehicle (N = 9) or R291-SG3199 at 0.15 mg/kg (N = 8), or 0.3 mg/kg (N = 8). Injection days are marked by arrows. B, Body weight curves of animals in A. C, Kaplan-Meier survival plot of the study in A. D, Tumor growth curves of SKNBE2 xenografts treated with vehicle (N = 7) or R291-SG3199 at 0.3 mg/kg (N =8). Drug injection days are marked by arrows. E, Body weight curves of animals in D. F, Kaplan-Meier survival plot of the study in D.

### RSPO2 furin mutant peptibody conjugated with CPT2 had potent and specific cytotoxic activity against cancer cells expressing LGR4/5 *in vitro* and *in vivo*

Given the lack of total inhibition of tumor growth by R291-SG3199 in NB xenograft models *in vivo* and the limitation of dosing levels of PBD-based ADCs due to the extremely high cytotoxic potency of free PBDs^50^, we reasoned that conjugation of RSPO2-based peptibodies with the camptothecin analog CPT2 may enhance efficacy without increasing toxicity. Previously, we generated a PDC (R462-CPT2) using an RSPO4-based peptibody called R462 with close to 8 drugs per peptibody^43^. To generate an RSPO2-based peptibody that is conjugated to CPT2, we applied a similar approach by replacing the N295A change of R291 and R299 with N295Q and created R290 and R296, respectively (Fig. 5A-C). Introduction of N295Q created two Gln residues in each Fc domain that can be conjugated with a bis-Azide linker to enable the conjugation of 8 drugs per peptibody^43^. R290 showed high affinity binding to LGR5, just like R291, whereas R296 had little affinity (Fig. 5D-F). R290 and R296 were then conjugated with bis-azide linker using microbial transglutaminase, followed by linking to DBCO-PEG8-PKG-CPT2 as described for R462 and R465^43^. The resulting PDCs, R290-CPT2 and R296-CPT2, were confirmed by mass spectrometry to have ∼8 molecules per peptibody (Fig. 5C). R290-CPT2 and R296-CPT2 were then tested in binding to HEK293T cells expressing LGR4, LGR5, or LGR6. Similar to the unconjugated R290 and R296, R290-CPT2 retained high affinity binding to all three receptors whereas R296-CPT2 showed low binding with reduced affinity (Fig. 5G-I). We then tested cytotoxic activity of R290-CPT2 and R296-CPT2 in cancer cell lines side-by-side. As shown in Figure. 6A-C, R290-CPT2 showed IC50 of 0.9 nM, 0.3 nM, and 0.5 nM in SKNAS, SKNBE2, and LOVO cells, respectively, whereas R296-CPT2 had IC50 of 24, 20, and 21 nM, respectively. These results suggest that R290-CPT2 has potent and specific cytotoxic activity. We also examined the cytotoxic effects of R290-CPT2 and R296-CPT2 in non-cancer cell line HEK293T cells. The two PDCs showed similar potency with IC50 = 20 nM which was similar to R296-CPT2 in cancer cell lines, suggesting that high potency of R290-CPT2 is cancer cells was due to expression of LGR4/5/6 (Fig. 6). Next, we tested R290-CPT2 in xenograft model of SKNAS cell *in vivo*. R290-CPT2 was given at 5.0 mg/kg, twice per week, similar to that of R291-SG3199, for a total of 6 doses. As shown in Figure. 6H, R290-CPT2 inhibited tumor growth by 77% (p = 0.005 vs vehicle on Day 14) with no major adverse effect and no significant body weight changes. (Fig. 6I). R290-CPT2 also extended survival significantly (median survival of 37 days vs 19 days for vehicle, p = 0.0001, log-rank test) (Fig. 6J). Overall, these results demonstrated that RSPO2 peptibody conjugated CPT2 has potent anti-tumor activity *in vitro* and *in vivo* without gross toxicity.

**Figure 5.**
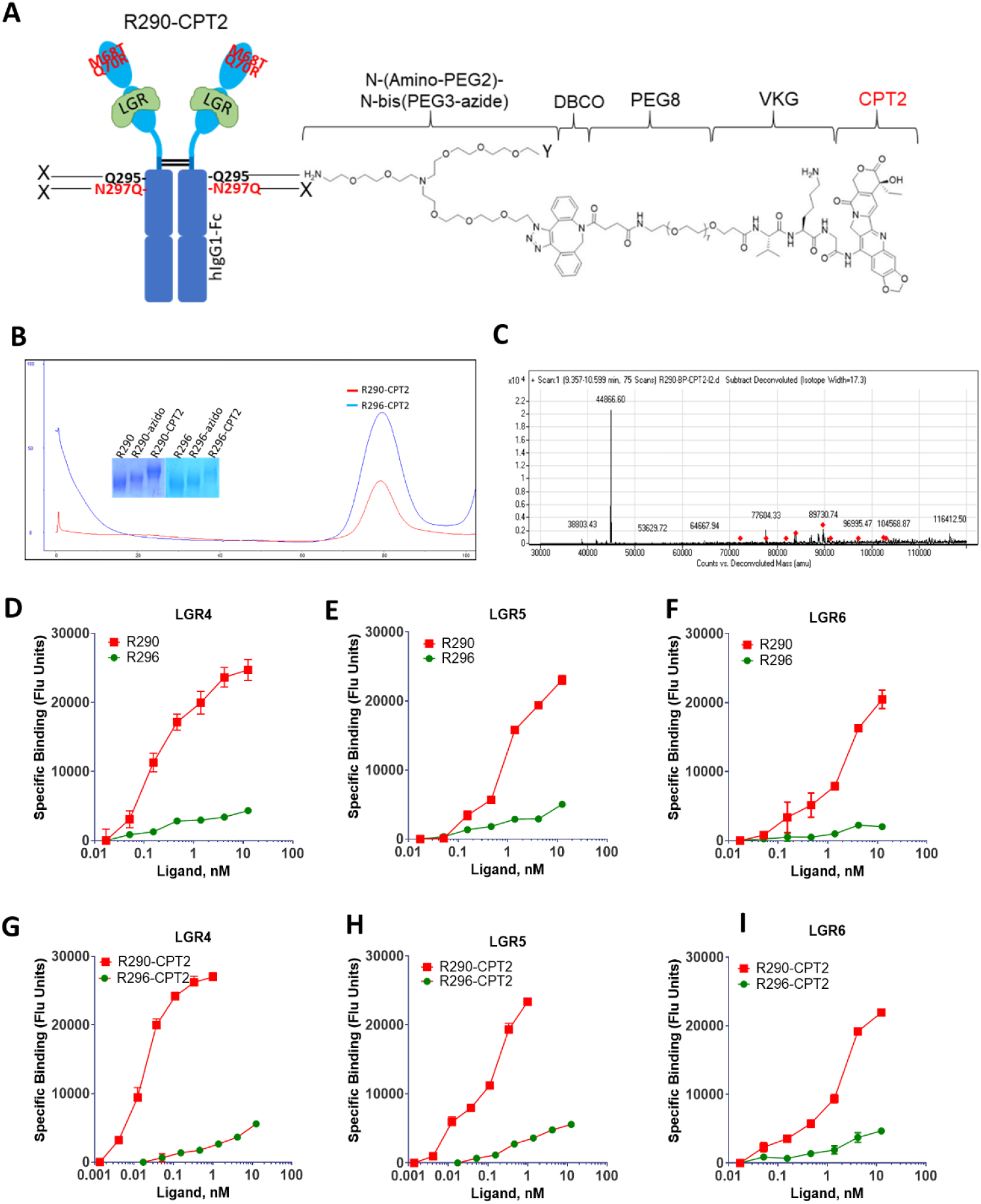
R290 and R296 PDC synthesis and conjugation. A, Schematic diagram of R290/R296 conjugated with CPT2. B, Chromatogram of R290- and R296-CPT2 in gel filtration and Coomassie staining of SDS-PAGE of R291/R299 before and after conjugation. C, Mass spectrum of R290-CPT2. D-F, Whole-cell binding of unconjugated R290 and R296 to HEK293T cells expressing LGR4 (D), LGR5 (E), and LGR6 (F). G-l, Whole-cell binding of R290-CPT2 and R296-CPT2 to HEK293T cells expressing LGR4 (G), LGR5 (H), and LGR6 (I). Error bars are SEM (N = 2-3).

**Figure 6.**
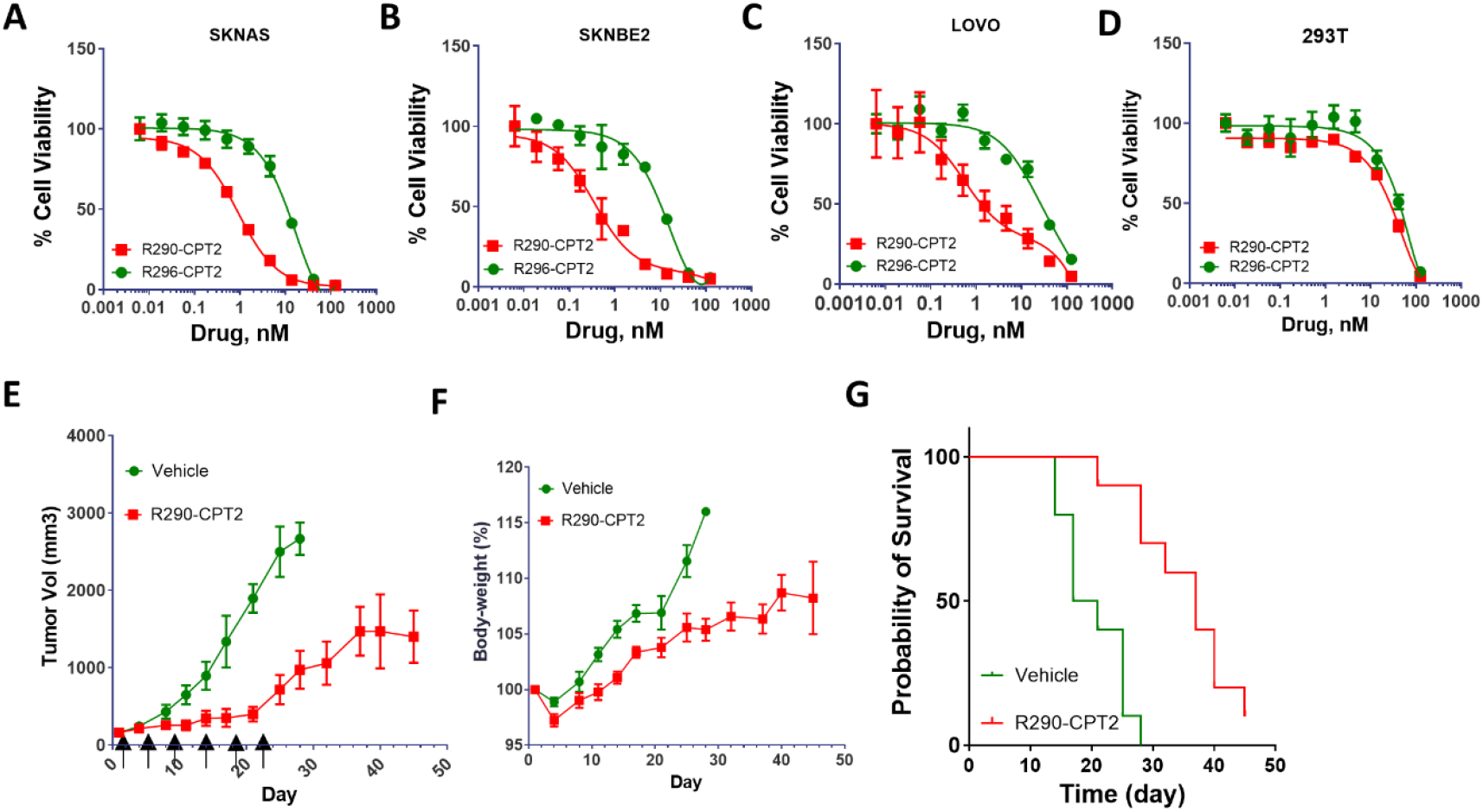
Activity of R290-CPT2 and R296-CPT2 in vitro and in vivo. A-C, Cytotoxic activity of R290-CPT2 and R296-CPT2 in SKNAS (A) and SKNBE2 (B), LOVO (C), and HEK-293T (D) cell lines. E, Tumor growth curves of SKNAS xenografts treated with vehicle or R290-CPT2 at 5 mg/kg (N = 10 per group). Injection days are marked by arrows. F, Body weight curves of animals in E. G, Kaplan-Meier survival plot of the study in E. Error bars are SEM (N = 2-3).

LGR4/5/6 are often co-expressed or alternately expressed at moderate to high levels in gastrointestinal cancers. In NB, LGR5 is highly expressed in high-risk tumors whereas LGR4 is expressed at moderate levels^37^. Simultaneous targeting all three receptors of LGR4/5/6 may offer increased efficacy and overcome tumor cell plasticity due to inter-conversion of LGR5-positive and -negative cells^26, 30^. We set out to develop a strategy that utilizes mutant ligands of LGR4/5/6 that retain high affinity receptor binding without potentiating Wnt/β-catenin signaling^43, 44^. Here, we generated new peptibodies based on a mutant form of RSPO2 furin domain and conjugated with either PBD or CPT-derived payloads. CPT-derived payloads are now the leading class for ADC development due to its success in solid tumors, and irinotecan/topotecan, both are derivatives of CPT, are widely used as chemotherapy for the treatment of CRC and high-risk NB^3, 51^. RSPO2 and RSPO4 are approximate 50% identical in the furin domain, yet RSPO2 is much more potent than RSPO4 in potentiating Wnt/β-catenin signaling as RSPO2 is able to inhibit RNF43/ZNRF3 in the absence of LGR4/5/6 ^46, 52^. We explored drug conjugation to an IgG1 that has a RSPO2 furin domain fused to its Fc for potentially longer *in vivo* stability and high affinity for LGR4^45^. Previously, we showed that a single mutation (Q70R) in the furin-1 domain of RSPO2 led to partial loss of activity in potentiating Wnt/β-catenin signaling^46^. As Zebisch et al showed that M68 of RSPO2-furin-1 was also important for its activity^48^, we generated a double mutant of RSPO2 (M68T/Q70R) and showed this mutation was completely inactive in potentiating Wnt/β-catenin signaling (Fig. 1D). The double mutation had no effect on binding affinity to LGR4/5/6, making it an suitable candidate for drug conjugation.

Conjugation of either PBD or CPT2 to RSPO2-based peptibodies achieved high potency *in vitro* in cancer cell lines expressing any of LGR4/5/6, yet the PDCs were not able to eradicate cancer after two weeks of treatment. Previously, we conjugated MMAE or CPT2 to RSPO4-based peptibodies, and both PDC were able to inhibit tumor growth completely or even induce tumor remission at high dose levels with CPT2-conjugated RPSO4 PDC^43, 44^. CPT2 conjugated to RSPO2- or RSPO4-based PDCs resulted in similar cytotoxic potency in vitro in the same cancer cell lines, consistent with the similar binding affinity of RSPO2 and RSPO4 to LGR4/5/6, especially when presented in the form of dimeric IgG1 Fc domain^53^. A major rationale in evaluating of RSPO2-based PDCs was the potential of longer half-life of RSPO2 peptibodies in the mouse based on early preliminary data. Unfortunately, RSPO2 and RSPO4 peptibodies had similar, relatively short half-live *in vivo*, at least in the mouse. When conjugated with two molecules of the PBD analog SG3199, R291-SG3199 had good solubility and showed high cytotoxic potency in cell lines expressing any of LGR4/5/6. This potency is higher than that of an anti-LGR5 antibody (8F2) conjugated with the same linker-payload^37^, suggesting that natural ligand-based drug conjugate may target LGR5 more effectively. However, 8F2-SG3199 and R291-SG3199 showed similar efficacy *in vivo* despite the much higher cytotoxic potency of R291-SG3199 *in vitro*. The most likely explanation for similar potency between 8F2-SG3199 and R291-SG3199 is that the half-life of R291-SG3199 is predicted to be much shorter than 8F2-SG3199 *in vivo*. Therefore, improving in vivo stability of RSPO2/RSPO4-peptibodies will be key to the development of RSPO2/RSPO4 PDCs for cancer treatment. Since the molecular weights of the peptibodies/PDCs are approximately 80 kDa, they are not cleared by the kidney, instead, most likely by protease-mediated cleavage in circulation. Current efforts are focused on delineating how the peptibodies are degraded and identifying protease cleavage sites.

Another major aspect affecting potency and efficacy of ADC/PDC are the selection of linker-payload and conjugation method^4^. In the present and previous studies ^43, 44^, we used the linker-payloads that were proven to be most successful in the development of ADCs, i.e., MMAE, CPT, and SG3199. MMAE belong to the class of microtubule inhibitors, and this class of payloads were proven to be mostly effective for hematological cancers and bladder cancer^4^. Microtubule inhibitors, however, are not effective for CRCs, which is the major reason we focused on CPT2 as payload for proof-of-principle. PBD payloads such as SG3199 are highly potent but often too toxic, making its ADCs only usable at low dosages which may limit tumor penetration^50, 54^. Given the success of deruxtecan (Dxd)-based ADCs for the treatment of solid tumors, various proprietary derivatives of CPT have now been generated and used for ADCs in different stages of clinical development^4, 55^. We selected CPT2 given its increased potency and solubility over Dxd^51^, and used the chemoenzymatic method for conjugation^49^. As RSPO furin domain contains five disulfide bonds that are critical for its structure^48^, direct conjugation of the inter-chain disulfide bonds of the IgG1-Fc domain of RSPO peptibodies is not feasible using TCEP (tris(2-carboxyethyl)phosphine)-based reduction that is used for conjugation of typical IgG1 antibodies. Future efforts will also develop strategies that will allow conjugation to the cysteine residues of the peptibodies.

## Conclusions

In conclusion, we generated two new peptibodies based on a mutant RSPO2 furin domain that showed high affinity, specific binding to LGR4/5/6 without potentiating Wnt/β-catenin signaling. The drug conjugate of the peptibodies displayed potent cytotoxic activity in LGR4/5/6-expressing CRC and NB cell lines *in vitro*, and robust anti-tumor efficacy *in vivo* without major adverse effect. Future studies will focus on optimizing the *in vivo* stability and potency of the peptibody by exploring alternative payload, Fc engineering strategies, and linker chemistry.

## EXPERIMENTAL SECTION

### Cell lines

The NB and CRC cell lines SKNAS, SKNBE2, CHP212, HT29, SW620, and RKO were purchased from the American Type Culture Collection (ATCC). CRISPR/Cas9-mediated gene editing via lentiviral vectors, as described previously^46, 56^, was utilized to generate LGR5-knockout variants of CHP212 cell lines. HEK292T cells stably expressing human LGR4, LGR5, or LGR6 and SuperTopFlash (STF) cell line were described in a previous study^11^.

### PDC preparation and characterization

The sequences encoding wild-type RSPO2 furin domain (R203) fused to human IgG1-Fc were described previously^46^. Its mutants (R291, R299, R290, and R296) were generated and cloned into the mammalian protein expression vector pCEP5 using the In-Fusion cloning method. To enable microbial transglutaminase (MTGase)-mediated site-specific conjugation, mutations were introduced at the N297 glycosylation site, generating N297A Fc variants for R291 and R299, and N297Q Fc variants for R290 and R296. Protein production and characterization were carried out as described before^46^. Briefly, peptibodies were expressed in Expi293™ cells (Thermo Fisher Scientific) by transient transfection, purified by CaptivA® HF Protein A Resin (Repligen) via gravity flow, and further refined through size-exclusion chromatography on a HiLoad 16/600 Superdex 200 pg column (Cytiva).

Subsequently, purified peptibodies (5 mg/mL in PBS, pH 7.2) were conjugated overnight at room temperature using 8% Activa® TI Transglutaminase (Ajinomoto) and 40 molar equivalents of amino-PEG4-azide linker (BroadPharm) or N-(Amino-PEG2)-N-bis(PEG3-Azide) linker (BroadPharm). Unreacted linker and residual MTGase were removed through Protein A resin treatment with gentle mixing at room temperature for 2 hours. The resulting conjugate was then reacted with 1.5 molar equivalents of linker-drug at room temperature for an additional 4 hours, provided as a 20 mg/m in dimethyl sulfoxide (DMSO) stock solution of either DBCO-PEG8-VA-PAB-SG3199 or DBCO-PEG8-Val-Lys-Gly-14-aminomethyl (CPT2) (Levena Biopharma), ensuring the final DMSO concentration remained below 10% (v/v). Following the reaction, excess reagents were removed by size-exclusion chromatography, and the final peptibody-drug conjugate (PDC) was buffer-exchanged into formulation buffer (20 mM sodium succinate, 6% trehalose, pH 5.0) stored at −80°C until further use.

PDC samples were analyzed by hydrophobic interaction chromatography on an Agilent 1260 Infinity II HPLC system equipped with an AdvanceBio HIC column (4.6 × 100 mm, 3.5 µm; Agilent). Samples (40 µL) were maintained at 10 °C prior to injection and separated at 25 °C using a flow rate of 0.8 mL/min. Mobile phase A consisted of 50 mM potassium phosphate and 1.5 M ammonium sulfate (pH 7.0), and mobile phase B contained 50 mM potassium phosphate with 10% isopropanol (pH 7.0). The gradient program was as follows: 0–3 min, 5% B; 3– 30 min, 5–100% B; followed by a 10 min post-run. Eluted species were monitored at 280 nm.

For intact mass analysis, PDCs were further characterized on an Agilent 6538 Q-TOF LC/MS coupled to an Agilent 1200 HPLC as described before^37^. Separation occurred on an Agilent PLRP-S reversed-phase column using a gradient of acetonitrile-water-formic acid solvents. Mass spectrometry was performed in positive ESI mode, and raw data were processed using Agilent MassHunter BioConfirm software for molecular mass determination.

### Cell based signaling and binding assay

Potentiation of Wnt/β-catenin signaling assay was performed using the SuperTopFlash cell line and 10% conditioned Wnt3A media as described before^11^. Whole-cell binding assays were conducted as previously described^11^. Briefly, stable HEK293-LGR4, HEK293-LGR5-DeltaC, and HEK293-LGR6 cells were seeded onto poly-D-lysine–coated black, clear-bottom 96-well plates and cultured overnight. Fc-tagged peptibodies and PDCs were serially diluted and added to the cells, followed by incubation on ice for 1 hour. Cells were then washed and fixed with 4.2% paraformaldehyde and incubated with anti-human Alexa Fluor 555 antibody. Fluorescence emission at 550 nm was measured using a plate reader (Tecan), and dose-response curves were fitted using GraphPad Prism (Boston MA) to calculate the half-maximum binding constant (Kd). All assays were performed at least three times, with duplicates or triplicates per experiment.

### Cytotoxic assay

For cytotoxicity assays, cells were seeded at a density of 3 × 10^3^ cells per well in 96-well plates (Costar Assay Plate, Corning). Serial dilutions of PDCs as indicated were then added. After incubation for 5 days, cell viability was assessed using the CellTiter-Glo® Luminescent Cell Viability Assay (Promega, Madison, WI) and measured on a TECAN Infinite® M1000 plate reader (Tecan Austria GmbH). The half-maximal inhibitory concentration (IC_50_) values were determined using GraphPad Prism software.

### Xenograft studies in mice

Animal studies were carried out in strict accordance with the recommendations of the Institutional Animal Care and Use Committee of the University of Texas at Houston (Protocol number AWC-19-0085 and AWC-21-0094). For SKNAS and SKNBE2 xenograft studies, female 9-10-week-old nu/nu mice (Charles River Laboratories) were subcutaneously inoculated with 5 × 10^6^ cells in 1:1 mixture with Matrigel (BD Biosciences). After 2 weeks, when tumor size reached approximately ∼100 mm^3^, mice were randomized and given vehicle (PBS), R291-SG3199 or R299-SG3199 at the indicated dose levels and dosing frequency by intraperitoneal injection. Mice were routinely monitored for morbidity and mortality.

Tumor volumes were measured approximately 2–3 times per week and estimated by the formula: tumor volume = (length × width^2^)/^2^.

### Statistical analysis

All data were analyzed using GraphPad Prism software. Data are expressed as mean ± SEM or SD as indicated in the Results section. For tumor volume analysis, one-way ANOVA with Dunnett’s multiple comparison test or Student’s unpaired two-tailed t-test for two-group comparisons was employed. Survival data were analyzed using Kaplan-Meier analysis with Log-rank (Mantel-Cox) test for P value calculation. P ≤ 0.05 was considered statistically significant.

## ASSOCIATED CONTENT

### Supporting Information

Supplementary Figures S1-3.

## AUTHOR INFORMATION

### Authors

**Yukimatsu Toh** – *Brown Foundation Institute of Molecular Medicine, University of Texas Health Science Center at Houston, Houston, TX 77030* USA;

**Jianghua Tu** – *Brown Foundation Institute of Molecular Medicine, University of Texas Health Science Center at Houston, Houston, TX 77030* USA

**Ling Wu** – *Brown Foundation Institute of Molecular Medicine, University of Texas Health Science Center at Houston, Houston, TX 77030* USA

**Adela M. Aldana** – *Brown Foundation Institute of Molecular Medicine, University of Texas Health Science Center at Houston, Houston, TX 77030* USA

**Jake J. Wen** – *Brown Foundation Institute of Molecular Medicine, University of Texas Health Science Center at Houston, Houston, TX 77030* USA

**Lynn H. Su** – *Brown Foundation Institute of Molecular Medicine, University of Texas Health Science Center at Houston, Houston, TX 77030* USA

**Li Li** – *Brown Foundation Institute of Molecular Medicine, University of Texas Health Science Center at Houston, Houston, TX 77030* USA

**Sheng Pan** – *Brown Foundation Institute of Molecular Medicine, University of Texas Health Science Center at Houston, Houston, TX 77030* USA

**Xiaowen Liang** – *Wntrix, K2Bio, 2710 Reed Rd*., *Suite 160, Houston, TX 77051 USA*.

**Jie Cui** – *Wntrix, K2Bio, 2710 Reed Rd*., *Suite 160, Houston, TX 77051 USA*

**Bin Yang** – *The Verna and Marrs McLean Department of Biochemistry and Molecular Pharmacology, Baylor College of Medicine, Houston, Texas 77030, USA, and Center for NextGen Therapeutics, Baylor College of Medicine, Houston, Texas 77030, USA*

**Jin Wang** – *The Verna and Marrs McLean Department of Biochemistry and Molecular Pharmacology, Baylor College of Medicine, Houston, Texas 77030, USA; Center for NextGen Therapeutics, Baylor College of Medicine, Houston, Texas 77030, USA, and Department of Molecular and Cellular Biology, Baylor College of Medicine, Houston, Texas 77030, USA*.

### Author Contributions

Conceptualization, Q.J.L. and Y.T.; methodology, Y.T., J.T., L.W., A.M.A., J.J.W., L.H.S., B.Y., X.L., L.L., S.P., J.W. and J.C.; writing—original draft manuscript, Y.T. and Q.J.L.; writing—review and editing, Y.T., L.H.S. and Q.J.L.; funding acquisition, Q.J.L. All authors have read and agreed to the published version of the manuscript.

### Notes

The authors declare the following competing financial interest(s): C.J., Q.J.L., and the Regents of the University of Texas have filed a provisional patent application related to this project. J.W. is the co-founder of Chemical Biology Probes LLC. J. W. has stock ownership in CoRegen Inc and serves as a consultant for this company. J.W., and B.Y. are the co-founders of Fortitude Biomedicines, Inc. and hold equity interest in this company. A version of the manuscript was deposited the preprint server bioRxiv

## ACKNOWLEDGEMENTS

This work was supported in part by funding from the Cancer Prevention and Research Institute of Texas (CPRIT) RP220169 (to Q.J.L.) and RP210119 (to Q.J.L.), from the National Cancer Institute of the National Institutes of Health under grant numbers R21CA267381 (to Q.J.L.) and R41CA281553 (to C.J. and Q.J.L), the Janice David Gordon for Bowel Cancer Research Endowment (to Q.J.L.), and Cancer Prevention & Research Institute of Texas (CPRIT, RP220480 to J.W.), the seed fund to Center for NextGen Therapeutics, and the Michael E. DeBakey, M.D., Professorship in Pharmacology (to J.W.).

## ABBREVIATIONS

LGR4/5/6: leucine-rich repeat containing, G protein-coupled receptors 4, 5, and 6
CRC: colorectal cancer
NB: neuroblastoma
RSPOs: R-spondins
PBD: pyrrolobenzodiazepine dimer
CPT2: camptothecin derivative
PDCs: peptibody-drug conjugates
ECD: extracellular domain
7TM: seven transmembrane
GI: gastrointestinal
TCGA’s: The Cancer Genome Atlas’s
ADCs: Antibody-drug conjugates
ATCC: American Type Culture Collection;

