## Supplementary Figures for "RSPO2-based peptibodies conjugated with pyrrolobenzodiazepine dimer or camptothecin analogs demonstrate potent anti-tumor activity by targeting the three receptors LGR4/5/6 in colorectal cancer and neuroblastoma"

### Table of Contents.

1) Figure S1. Cytotoxicity of Free Payloads in Neuroblastoma Cell Lines.
2) Figure S2. Pharmacokinetic profiles of R291 and R291-SG3199 in mice
3) Figure S3. Hydrophobic Interaction Chromatography (HIC) Analysis of Peptibody–Drug Conjugates

**Figure S1.** Cytotoxicity of Free Payloads in Neuroblastoma Cell Lines. Dose–response cytotoxicity of free monomethyl auristatin E (MMAE), pyrrolobenzodiazepine dimer (PBD), deruxtecan (DXd), and SN38 was evaluated in neuroblastoma cell lines using a 5 days viability assay. Curves and corresponding $IC_{50}$ values were generated by nonlinear regression. Panels show (A) SK-N-AS, (B) SK-N-BE(2), (C) CHP-212, and (D) CHP212-LGR5KO cells. Data represent mean ± SEM from independent experiments.

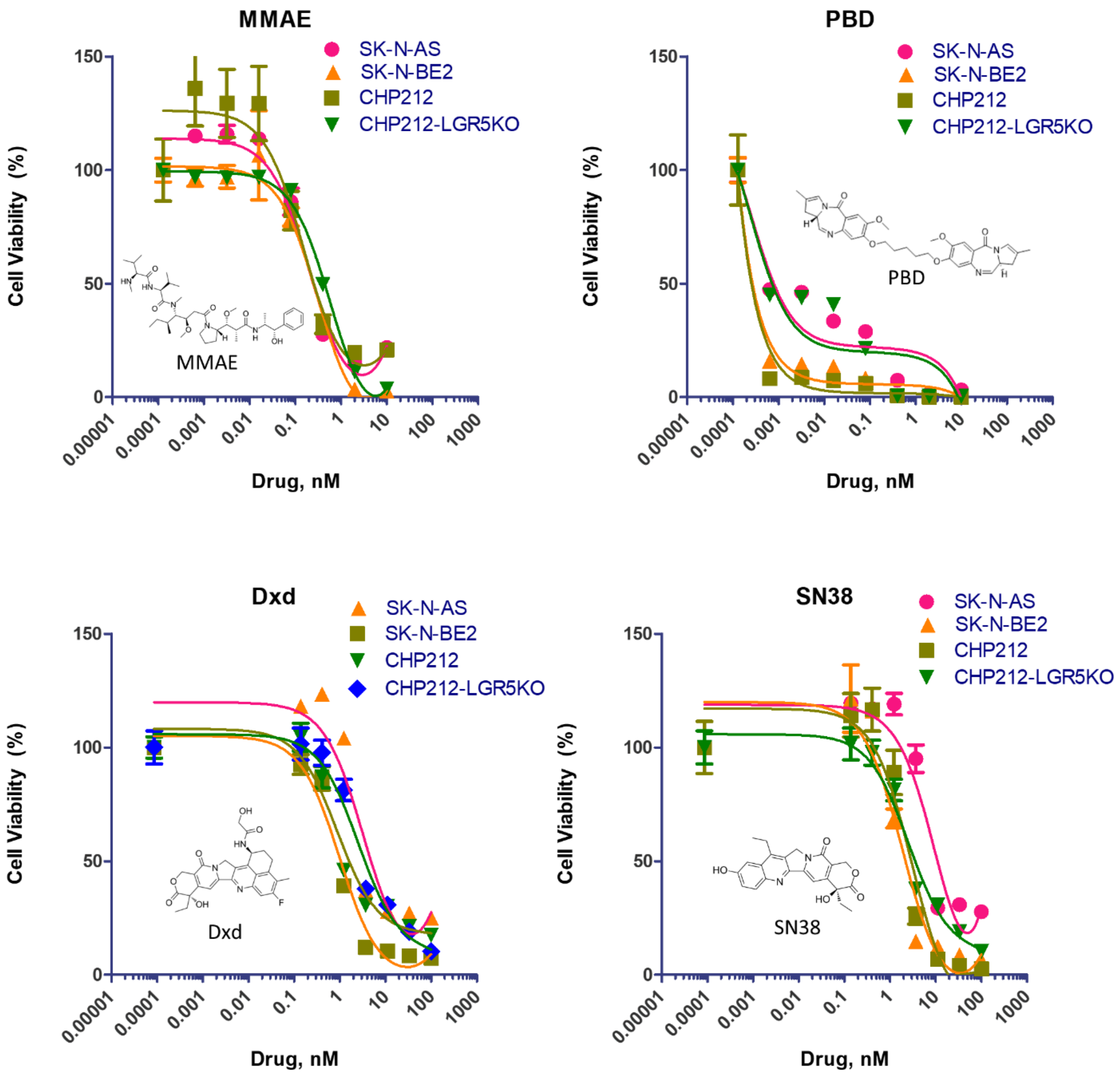

Figure S2. Pharmacokinetic profiles of R291 and R291-SG3199 in mice. Analysis was performed in C57BL/6 mice (n = 3 per group) following a single intravenous dose.

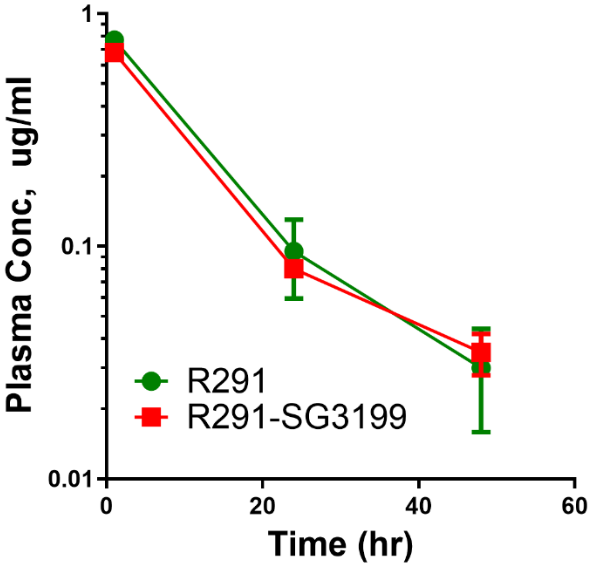

Figure S3. Hydrophobic Interaction Chromatography (HIC) Analysis of Peptibody–Drug Conjugates. HIC profiles of purified peptibody and peptibody–drug conjugates (PDCs) were generated to assess conjugation quality, hydrophobicity changes, and drug-to-antibody ratio (DAR) distribution. Chromatograms illustrate the shift in retention characteristics following payload conjugation compared with the unconjugated peptibody.

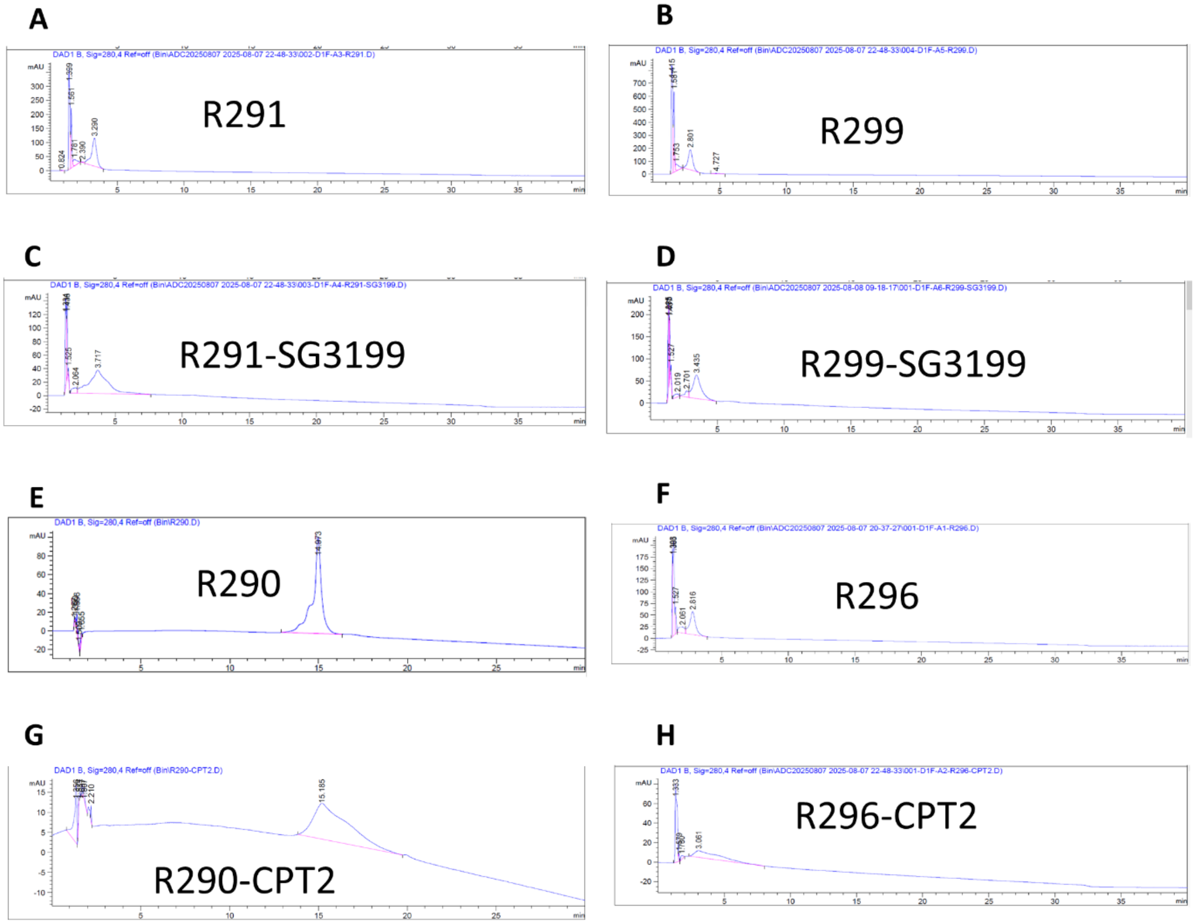
